# Comparative analysis of IgG Responses to recombinant Qβ phage displayed MSP3 and UB05 in Dual HIV-malaria infected adults living in areas differing in Malaria transmission intensities

**DOI:** 10.1101/303628

**Authors:** Abel Lissom, Herve F. Ouambo, Rosette Megnekou, Malachy I. Okeke, Loveline N. Ngu, Palmer M. Netongo, Apeh A. Ngoh, Carrie A. Sanders, Swapnil Bawage, Thibau F. Tchouangueu, Colince J. Tchadji, Arinze S. Okoli, Ghislain D. Njambe Priso, Rosario Garcia, Anna Gutiérrez, George O. Chukwuma, Charles O. Esimone, Eric A. Achidi, Wilfred F. Mbacham, Lazare Kaptue, Rose FG Leke, Chae Gyu Park, Alain Bopda Waffo, Godwin W. Nchinda

## Abstract

Immunoglobulin G specific responses against *Plasmodium falciparum* merozoite antigens such as the merozoite surface protein 3 (MSP3) and UB05 are known to play critical roles in parasitemia control and protection from symptomatic illness. However when there is intense perennial malaria transmission coupled with concurrent infection with the human immunodeficiency virus type 1 (HIV), knowledge of IgG antibody response profiles is limited. In this study we assessed the impact of dual HIV-Malaria infections on IgG subclass responses to MSP3 (QβMSP3) and UB05 (QβUB05) in individuals living in two areas of Cameroon differing in transmission intensity. We observed differences in antigen specific IgG and IgG subclass responses which was dependent upon the antigen type, malaria transmission intensity, HIV infection, malaria infection and dual HIV-malaria infections. Individuals living in high malaria transmission areas irrespective of HIV or malaria status had significantly higher IgG responses to both antigens (P=0.0001 for QβMSP3, P=0.0001 for QβUB05) than their counterpart from low transmission areas. When dual HIV-Malaria infection is considered significantly higher QβMSP3 specific IgG1 (P=0.0001) and IgG3 (P=0.04) responses in double negative individuals was associated with protection against malaria in low transmission areas. Superior QβUBO5 specific IgG1 responses (P=0.0001) in double negative individuals were associated with protection in high transmission areas in contrast to significantly higher IgG3 responses to QβUB05 (P=0.0001) which were more relevant to protection in low malaria transmission areas in the same population. Thus, understanding immune responses to QβUB05 and QβMSP3 could facilitate the development of immunotherapeutic strategies suitable for areas differing in malaria transmission intensity.

## Introduction

In Cameroon like in most sub Saharan African countries people living in low or high malaria transmission areas are exposed to different frequencies of *Plasmodium falciparum*. Within such regions long term inhabitants suffer repeated exposure to varying strains of malaria parasite during the course of several years eventually developing protection from infection and symptomatic illness irrespective of malaria transmission intensity (1). A critical component of this naturally acquired immunity is *Plasmodium falciparum* induced IgG and IgG subclass antibody responses targeting a number of parasite derived antigens. However little is known about the IgG antibody subclass profile that mediates protective immunity to malaria in both low and high malaria transmission areas. Also, when there is concurrent infection with the human Immunodeficiency virus type 1(HIV), knowledge of the precise nature of parasite antigen directed IgG subclass antibody responses functional in the afflicted individuals is limited. Given that HIV infection depletes the immune system there is need to understand the role of promising malaria target antigens and the profile of IgG subclass responses driving protective immunity to malaria in both low and high transmission areas.

Antibodies to several asexual blood stage antigens including apical membrane antigen 1 (AMA-1), erythrocyte binding antigen (EBA-175), the merozoite surface proteins (MSPs), reticulocyte-binding protein homologue (Rh5), Glutamate-rich protein (GLURP), UB05 and circumsporozoite protein (CSP) have been demonstrated to be an essential component of naturally acquired immunity reducing parasite multiplication thereby preventing infection and clinical disease in long term inhabitants of endemic regions (2–8). In this regards high levels of IgG antibody subclass responses and diversity of the target antigens have been associated with naturally acquired immunity to malaria (6–9). However, due to inherent polymorphism in the asexual blood stage antigens (10) some elements of antibody-mediated immunity to *P. falciparum* have been reported to be strain specific (11, 12) thereby limiting their utility as global malaria vaccine candidates.

In low and high transmission areas the attainment of clinical immunity against malaria or protection from infection is largely dependent upon continuous exposure to multiple parasite variants (13) leading to an accumulation of a broad range of antibody specificities responsible for the naturally acquired immunity. In areas differing in transmission intensities it is uncertain which IgG subclass respond profiles are relevant to naturally acquired immunity. The scenario becomes even more challenging when there is attendant co-infection with HIV which dysregulates antibody responses and depletes the immune system.

In this study we have determined in low and high malaria transmission areas the impact of dual HIV-malaria infection on IgG subclass responses to two conserved *P. falciparum* derived asexual blood stage antigens displayed separately upon a recombinant RNA coliphage Qβ as previously described by our group (14, 15). The recombinant phage QβMSP3 displays the conserved C-terminal 88 aa of the merozoite surface protein 3 (16, 17) whilst QβUB05 bears the previously described malaria antigen UBO5 (18). Surface display upon the recombinant RNA coliphage Qβ as previously demonstrated by our group improves the antigenicity of inserted antigens (14, 15).

Antibodies specific to *Plasmodium falciparum* MSP3 are known to mediate parasite killing in association with monocytes in a process referred to as (19–21) antibody-dependent cellular inhibition (ADCI). MSP3 specific antibodies therefore contribute in preventing symptomatic disease through the inhibition of blood parasite invasion cycles ultimately leading to a reduction in parasite burden and episodes of malaria (22, 23). A number of malaria vaccine candidates incorporating this highly conserved C-terminal end of MSP3 have been assessed in clinical trials with promising outcomes (19, 24–26). On other hand UB05 specific antibodies have also been associated with protection in exposed populations (18). We compared between dual HIV-malaria infected and double negative individuals the IgG subclass responses specific to the malaria vaccine antigens in both low and high malaria transmission area of Cameroon. Our study can facilitate the identification of surrogate markers of malaria immunity useful in the design of novel highly efficacious vaccines and the development of immunotherapeutic strategies to enhance immunity to malaria in people living in areas differing in transmission intensity.

## Materials and Methods

### Study site

The study was carried out in two areas of Cameroon (Yaounde and Bikop) differing in malaria transmission intensity. As the Capital city of Cameroon Yaounde (3°52ʹN11°31ʹE) is a multiethnic city situated at an average elevation of 750 m. Bikop on the other hand is a remote rural area located 48 KM away from Yaounde with year round intense malaria transmission. Both Yaoundé and Bikop are holendemic for malaria however with differing transmission intensity. This is mainly because unlike Yaounde, Bikop is located in the heart of the rain forest with a large number of mosquitoe breedings sites and poorly constructed houses favoring sustained high malaria transmission. The temperature in both areas is around 23.7°C with a similar average annual rain fall of 1643 mm. There also have similar rainy (March to June, September to November) and dry seasons (December to February, July–August) (27).

### Ethical clearance

This study received ethical approval from the Cameroon National Ethics Committee for Human Health Research (Reference numbers 2015/03/561/CE/CNERSH/SP and 2018/01969/CE/CNERSH/SP) and the CIRCB institutional review board (protocol number 14-11). All participants provided written informed consent. Data were processed using specific identifiers for privacy and confidentiality purposes. Clinical data generated during the course of this study was provided free of charge to all participants.

### Study design

This was a cross-sectional study which enrolled HIV-1 infected and non-infected people who were 21 years or older. Participants with other infection (including microfilaria, dengue, TB, and hepatitis B and C) and pregnant women were excluded from the study. All participants were members of the CIRCB AFRODEC cohort (28–30)

### Study area

This study was carried out in two areas of Cameroon (Yaounde and Bikop) differing in malaria transmission intensity. As the Capital city of Cameroon Yaounde (3°52ʹN11°31ʹE) is a multiethnicity situated at an average elevation of 750 m. Bikop is malaria hotspot with a high incidence of malaria located 48 KM away from Yaounde with year round intense malaria transmission. Both Yaoundé and Bikop are holoendemic for malaria however with differing transmission intensities. This is mainly because unlike Yaounde, Bikop is located in the heart of the rain forest with a high density of mosquitoes and poorly constructed houses favoring sustained high malaria transmission. The temperature in both areas is around 23.7°C with similar average annual rain fall of 1643 mm. There also have similar rainy (March to June, September to November) and dry seasons (December to February, July–August).

### Study Population

A total of 124 participants were recruited for this study. All participants were adults participants of the CIRCB AFRODEC cohort 23-25. Participants were constituted in for groups consisting of dual HIV-malaria infected (HIV+/Mal+), HIV mono-infected (HIV+/Mal-), malaria monoinfected (HIV-/Mal+) and double negative (HIV-/Mal-) people.

### Plasma sample collection and Processing

About 4 ml of blood was collected into plastic Vacuum blood spray-coated K2EDTA tubes called Vacutest (Vacutestkirma, Italy). Subsequently, samples were transported to the Vaccinology laboratory of Chantal BIYA International Reference Centre (CIRCB) for storage and analysis. All samples were stored at room temperature and processed within 4 hours of collection. To obtain plasma, samples were centrifuged at 2,000 rpm for 10 min at 4°C. The plasma fraction was harvested sterile under the hood, aliquoted in small single-use volumes and stored at −20°C until use. The plasma obtained from participants was heat inactivated for 30 minutes at 56°C prior to ELISA assay.

### HIV infection and CD4 T cell Enumeration

Confirmation of HIV status was done as described for the CIRCB AFRODEC cohort using the Cameroon’s national algorithm for the diagnosis of HIV infection as previously reported for the CIRCB AFRODEC cohort (31)

Absolute numbers of helper CD4+ T cells for HIV+ participants were determined in fresh whole blood using BD multitest CD3/CD8/CD45/CD4 and TruCount tubes (BD biosciences, USA) according to the manufacturer’s instructions.

### Malaria Diagnosis and microscopy

A malaria rapid diagnostic test was done on the blood samples according to the manufacturer’s instructions (SD Bioline, USA). In addition, thick peripheral blood films were stained with Giemsa and examined using a microscope following standard quality-controlled procedures, for the presence of malaria parasites.

### Study antigens

The antigens consisted of recombinant Qβ displaying Plasmodium falciparum 3D7 strain sequence derived C-terminal part of MSP3 (QβMSP3) and UB05 (QβUB05) generated in our group as previously described (14, 15).

### Determination of IgG and IgG subclasses antibody responses specific to QβUB05 and QβMSP3

The plasma levels of antibodies specific to the malaria antigens QβUB05 and QβMSP3 were determined through ELISA assay. Briefly high binding ELISA plates were coated with 10^7^ particles/well of each recombinant phage and incubated overnight at 4°C. The following day, Plates were washed 3× with PBST (PBS containing 0.05% Tween 20) and blocked with 3% BSA in PBS for one hour at 37 °C. Heat inactivated plasma samples were diluted in PBS at 1:300 (for IgG detection) or 1:100 (for IgG subclasses detection), then 100 μl/well added in triplicate and incubated for two hours at 37 °C. The plates were washed four times with PBST after which the bound antibody was probed with the peroxidase-conjugated mouse anti-human IgG and IgG subclasses (IgG1, IgG2, IgG3 and IgG4) diluted 1:4000 in 1X PBS. Bound conjugate was detected using ABTS substrate and stop solution according to the manufacturer’s protocol (southern biotech, Birmingham USA). The colorimetric signal was measured at 405 nm using a multiscan FC microplate reader (Thermo Fisher Scientific, USA).

### Statistical analysis

Data analysis was performed with Graphpad Prism Software version 6.1. The data were expressed as median (25th percentile-75th percentile). Comparisons of medians among two groups were performed by the U-Mann-Whitney test. Statistical significance was confirmed when P < 0.05.

## Results

### High malaria transmission intensity is associated with superior QβMSP3 and QβUB05 specific IgG antibodies

Individuals in areas of high perennial malaria transmission intensity developed significantly higher (P=0.0001) levels of QβMSP3 and QβUB05 specific IgG responses than those living in low transmission zones (compare Fig 1A with B). In high malaria transmission regions QβMSP-3 specific IgG responses were comparatively higher but not significant different than responses specific to QβUB05 in both positive (P=0.09) and negative (P=0.08) individuals (Fig. S1). In addition no significant difference is observed in dual HIV-Malaria infected people in their plasma reactivity with the two recombinant antigens both in low and high transmission areas (Fig 1A&C). Similarly when people negative for both malaria and HIV (double negative individuals) were considered no significant difference is observed in the IgG responses specific to the two malaria vaccine antigens. The effect of HIV-1 infection was a significant reduction in IgG responses specific to the two antigens in both low (P=0.0001 for QβMSP3, P=0.04 for QβUB05) and high (P=0.0001 for QβMSP3) malaria transmission areas. Surprisingly, there was no difference in these values for UB05 in the high malaria transmission region (Fig. 1C&D).

**Figure 1:**
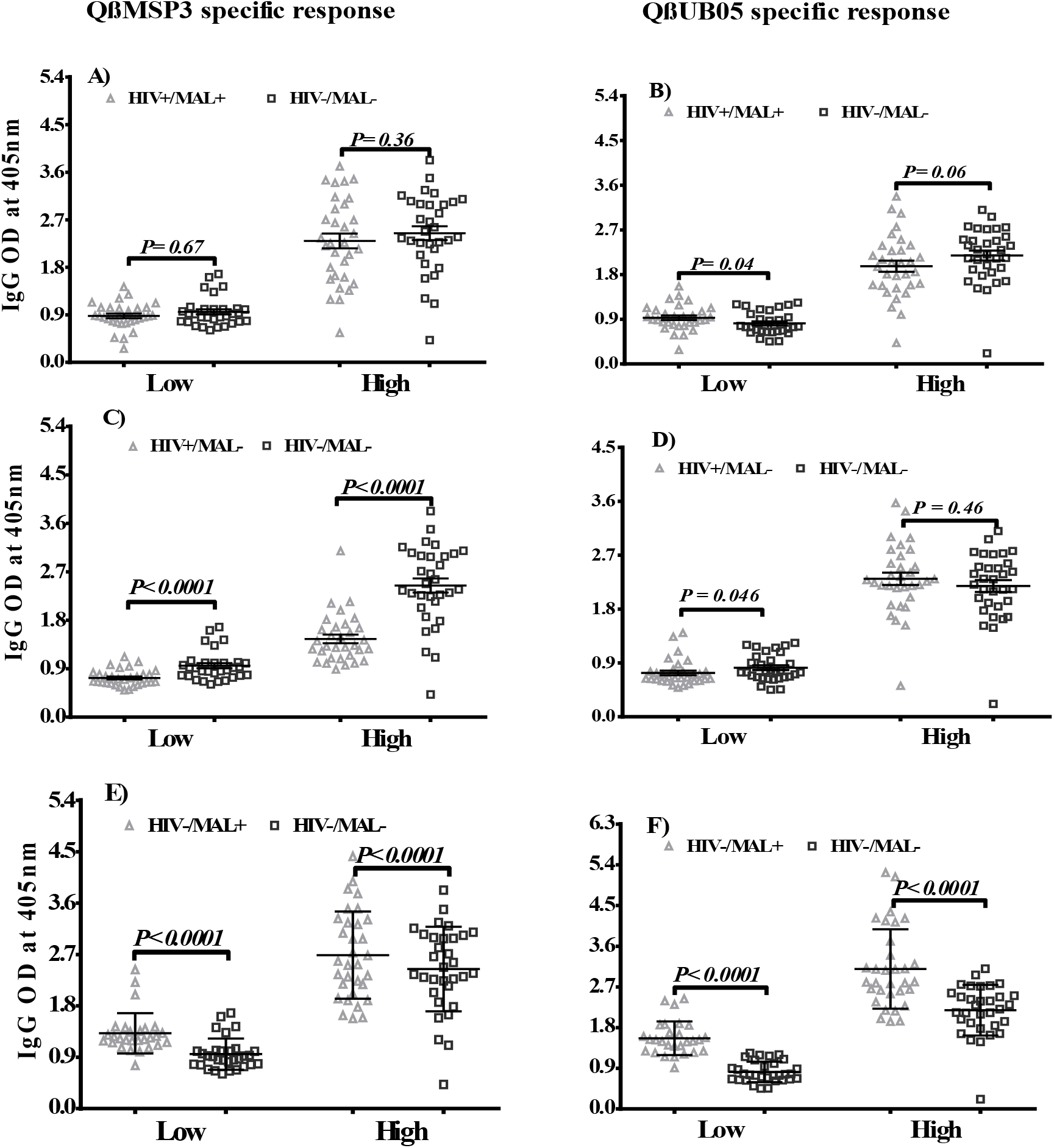
specific IgG responses to recombinant phages QβMSP3 and QβUB05 with respect to dual HIV-Malaria infections in individuals living in low and high malaria transmission areas. Comparison of IgG antibody responses between dual HIV-malaria infected and double negative individuals in low and high malaria transmission areas. IgG specific responses to the recombinant phages QβMSP3 **(A)** and QβUB05 **(B)** in low and high transmission areas. Effect of HIV infection on QβMSP3 **(C)** and QβUB05 **(D)** specific IgG responses in the two transmission areas. IgG responses specific to QβMSP3 (E) and QβUB05 (F) in individuals positive for *Plasmodium falciparum* in both transmission areas.

On the other hand, the overall effect of *Plasmodium falciparum* infection in both low and high malaria transmission regions was a significant increase in IgG responses specific to both antigens irrespective of dual HIV-malaria infection (Fig. 1E&F). Thus whereas HIV infection resulted to a significant a reduction in antigen specific IgG antibody levels *Plasmodium falciparum* infection resulted into a significant increase in the IgG antibody levels.

### IgGl subclass response in relation to Dual HIV-malaria infection and the intensity of malaria transmission

Whereas double negative individuals in low malaria transmission areas showed significantly higher (P=0.0001) IgG1 responses specific to QβMSP3 than to QβUB05; in high malaria transmission areas the converse was true (Fig. 2B) with IgG1 responses specific to QβUB05 being superior (P=0.0001). The effect of dual HIV-malaria infection in low transmission area was a significant reduction in the QβMSP3 specific IgG1 responses (P=0.0001) in contrast to individuals living in high malaria transmission areas where QβMSP3 specific IgG1 responses were similar to those of the double negative participants. On the other hand IgG1 responses specific to QβUB05 in high transmission areas remain comparatively higher than those to QβMSP3 in both dual HIV-malaria positive (P=0.004) and double negative (P=0.0001) individuals (Fig. 2A&B). This probably indicates the relevance of QβUB05 specific IgG1 in predicting malaria immunity in high transmission areas even under the challenging circumstances of HIV infection. The impact of dual HIV-malaria infection in low transmission areas was therefore a significant reduction (P=0.0001) in IgG1 responses specific to QβMSP3 (Fig 1C) in contrast to QβUB05 where IgG1 responses in this group remain higher than the double negative participants (P=0.0001). Again in a high transmission area the IgG1 responses to the two antigens were comparatively higher than values for the low transmission area. In addition IgG1 responses to QβUB05 in double negative individuals were also significantly higher (P=0.004) than dual HIV-malaria infected people. Thus in addition to malaria transmission intensity differences in IgG1 subclass antibody levels might also be dependent upon co-infection with HIV-1 and the antigen of choice.

**Figure 2:**
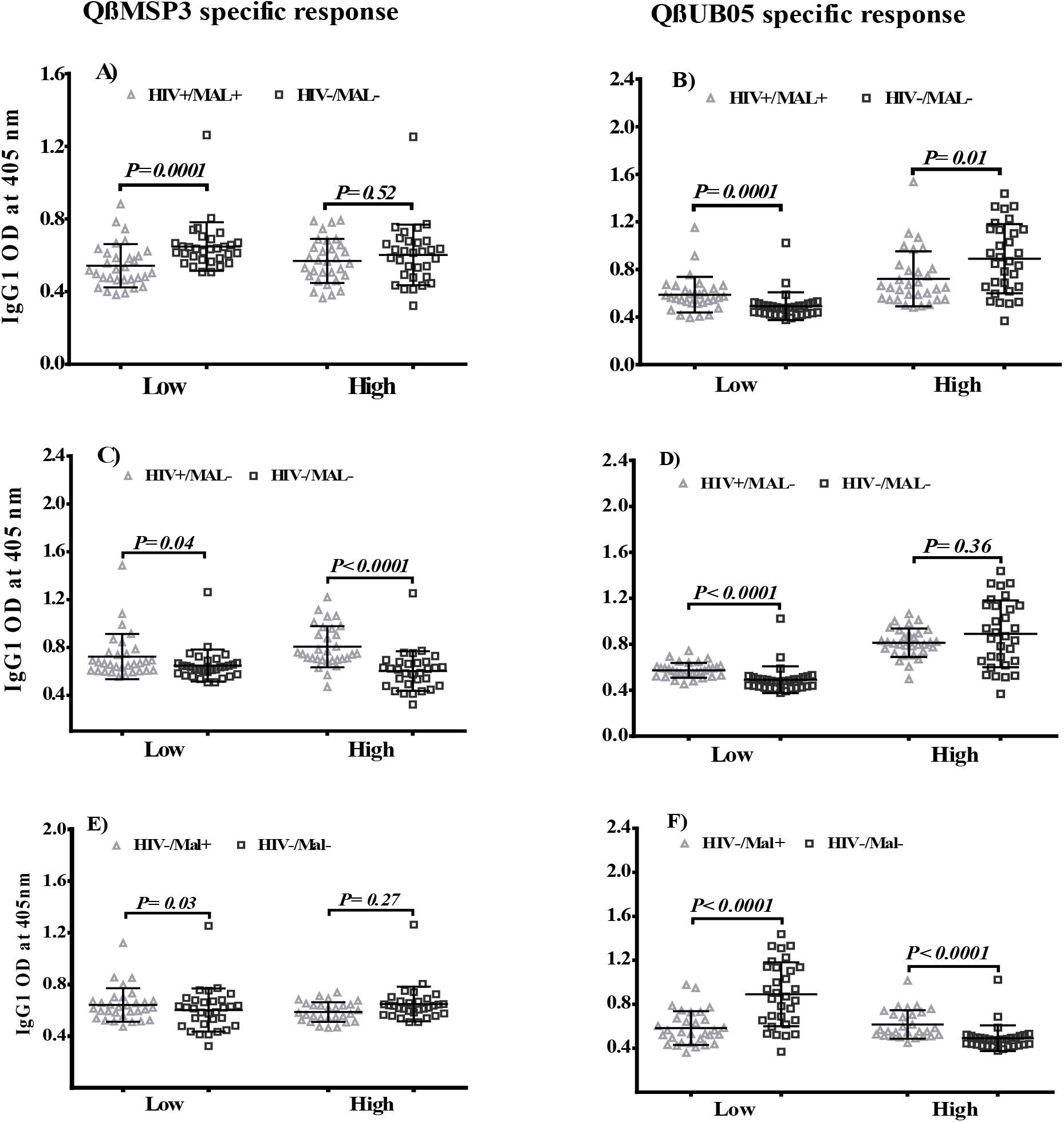
IgG1 specific response to recombinant phages QβMSP3 and QβUB05 with to dual HIV-Malaria infections in individuals living in low and high malaria transmission areas. Comparison of IgG1 antibody responses between dual HIV-malaria infected and double negative individuals in low and high malaria transmission areas. IgG1 responses specific to the recombinant phages QβMSP3 (A) and QβUB05 (B) in low and high transmission areas. Effect of HIV infection on QβMSP3 (C) and QβUB05 (D) specific IgG1 responses in the two transmission areas. IgG1 responses specific to QβMSP3 (E) and QβUB05 (F) in individuals positive for *Plasmodium falciparum* in both transmission areas.

Thus in a low malaria transmission area, significantly high IgG1 responses specific to QβMSP3 is associated with resistance to malaria in contrast to superior QβUB05 specific IgG1 antibodies which were instead relevant to protection in high a transmission area.

### IgG2 subclass response in relation to Dual HIV-malaria infection and the intensity of malaria transmission

In low malaria transmission areas significantly higher IgG2 subclass responses (P=0.0001 for QβMSP3 and P=0.0001 for QβUB05) were observed in double negative individuals relative to dual HIV-malaria positive participants (Fig. 3A&B). The converse was the case for high malaria transmission areas where dual HIV-malaria infection resulted to significantly higher IgG2 responses to both QβMSP3 (P=0.0001) and QβUB05 (P=0.0001). Overall the IgG2 responses specific to both antigen was comparatively superior in low relative to high transmission areas (Fig. 3A&B). The effect of an infection with HIV was a general increase in IgG2 responses specific to both antigens in low and high transmission areas (Fig. 3A&B). Several studies from other endemic areas of Africa have indicated that high circulating levels of malaria parasite antigen specific IgG2 could be a marker of severity of malaria infection (32–34). Except for MSP3 specific IgG2 an infection with plasmodium falciparum was associated with an increased in IgG2 responses to both antigens in all transmission areas. In a low malaria transmission area the impact of dual HIV-malaria infection was therefore a significant reduction in IgG2 responses to both antigens. In contrast in a high malaria transmission area dual HIV-malaria infection resulted to superior IgG2 responses to both QβMSP3 (P=0.0001) and QβUB05 (p=0.0001) relative to the double negative participants. Thus with respect to both antigens there is a differential IgG2 response between high and low malaria transmission areas.

**Figure 3:**
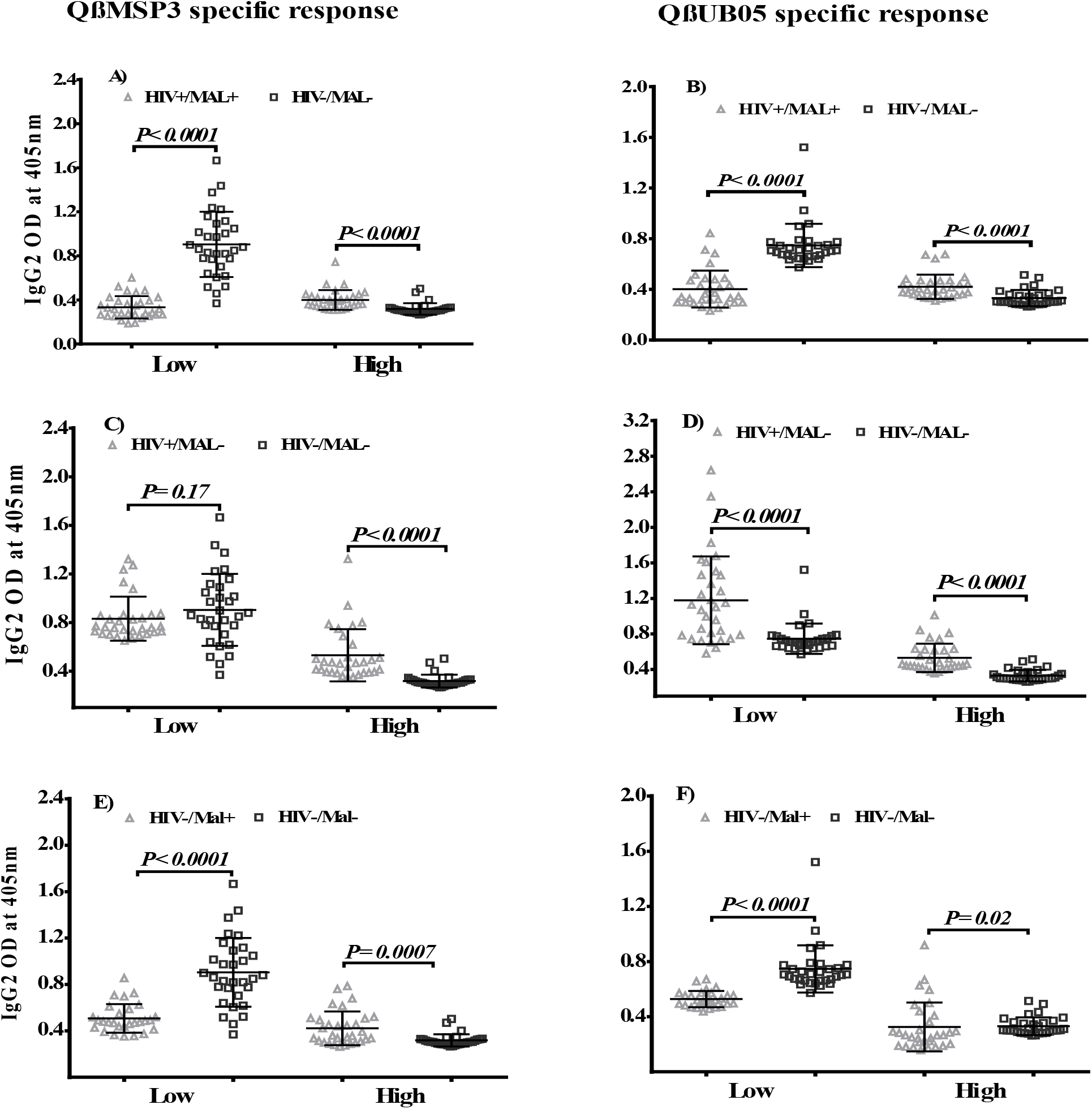
IgG2 responses specific to the recombinant phages QβMSP3 and QβUB05 with respect to dual HIV-Malaria infection in individuals living in low and high malaria transmission areas. Comparison of IgG2 antibody responses between dual HIV-malaria infected and double negative individuals in low and high malaria transmission areas. IgG2 responses specific to the recombinant phages QβMSP3 (A) and QβUB05 (B) in low and high transmission areas. Effect of HIV infection on QβMSP3 (C) and QβUB05 (D) specific IgG2 responses in the two transmission areas. IgG2 responses specific to QβMSP3 (E) and QβUB05 (F) in individuals positive for *Plasmodium falciparum* in both transmission areas.

### IgG3 subclass response in relation to Dual HIV-malaria infection and the intensity of malaria transmission

In low malaria transmission areas the IgG3 subclass antibody responses were significantly higher in double negative (p=0.0001 for QβMSP3 and p=0.0001 for QβUB05) compared to dual HIV-malaria infected participants. On the other hand in high malaria transmission areas no difference is observed between dual HIV-malaria infected and double negative individuals with respect to IgG3 subclass responses to both antigens. The effect of HIV infection was a significant reduction of IgG3 responses specific to QβMSP3 in both low (p=0.0003) and high (p=0.0001) malaria transmission areas. Similarly IgG3 specific responses to QβUB05 were significantly reduced in low (P=0.01) and high (P=0.0001) transmission areas after HIV infection. In low transmission areas *Plasmodium falciparum* infection resulted to a significant increase in IgG3 responses specific to both QβMSP3 (P=0.0003) and QβUB05 (P=0.0004) respectively. In contrast in high malaria transmission we observed significantly lower IgG3 responses specific to QβMSP3 (P=0.0001) and QβUB05 (P=0.0001) after Plasmodium falciparum infection. Thus there is differential effect of HIV and *Plasmodium falciparum* infections on IgG3 specific responses to both antigens which was also associated with the malaria transmission intensity.

### IgG4 subclass response in relation to Dual HIV-malaria infection and the intensity of malaria transmission

There was a differential expression of antigen specific IgG4 subclass responses between low and high malaria transmission areas (Fig. 5A&B). Overall for both antigens in low compared to high transmission areas dual HIV-malaria positive and double negative individuals showed significantly higher IgG4 responses. However in low transmission areas whereas QβMSP3 specific IgG4 responses in dual HIV-malaria infected individuals were superior to those of double negative individual (P=0.0001); for QβUB05 IgG4 responses the converse was true. On the other hand in high transmission areas no difference was observed between dual HIV-malaria infected and double negative individuals with respect to antigen specific IgG4 responses. The effect of HIV infection in all the malaria transmission areas was a significant reduction in IgG4 responses specific to both antigens (Fig.5 C&D). Plasmodium falciparum infection in low transmission areas resulted to a significant increase in IgG4 responses specific to QβMSP3 (P=0.0001) and QβUB05 (P=0.0001) respectively (Fig.5E&F). In contrast in high transmission areas the converse is true as IgG4 responses specific to both QβMSP3 (P=0.0001) and QβUB05 (P=0.0001) decreased significantly after Plasmodium falciparum infection (compare Fig. 5E with F). Thus variation in IgG4 specific responses was dependent upon several factors including transmission intensity, malaria parasite antigen, HIV infection, *Plasmodium falciparum* infection and dual HIV-malaria infection.

**Figure 4:**
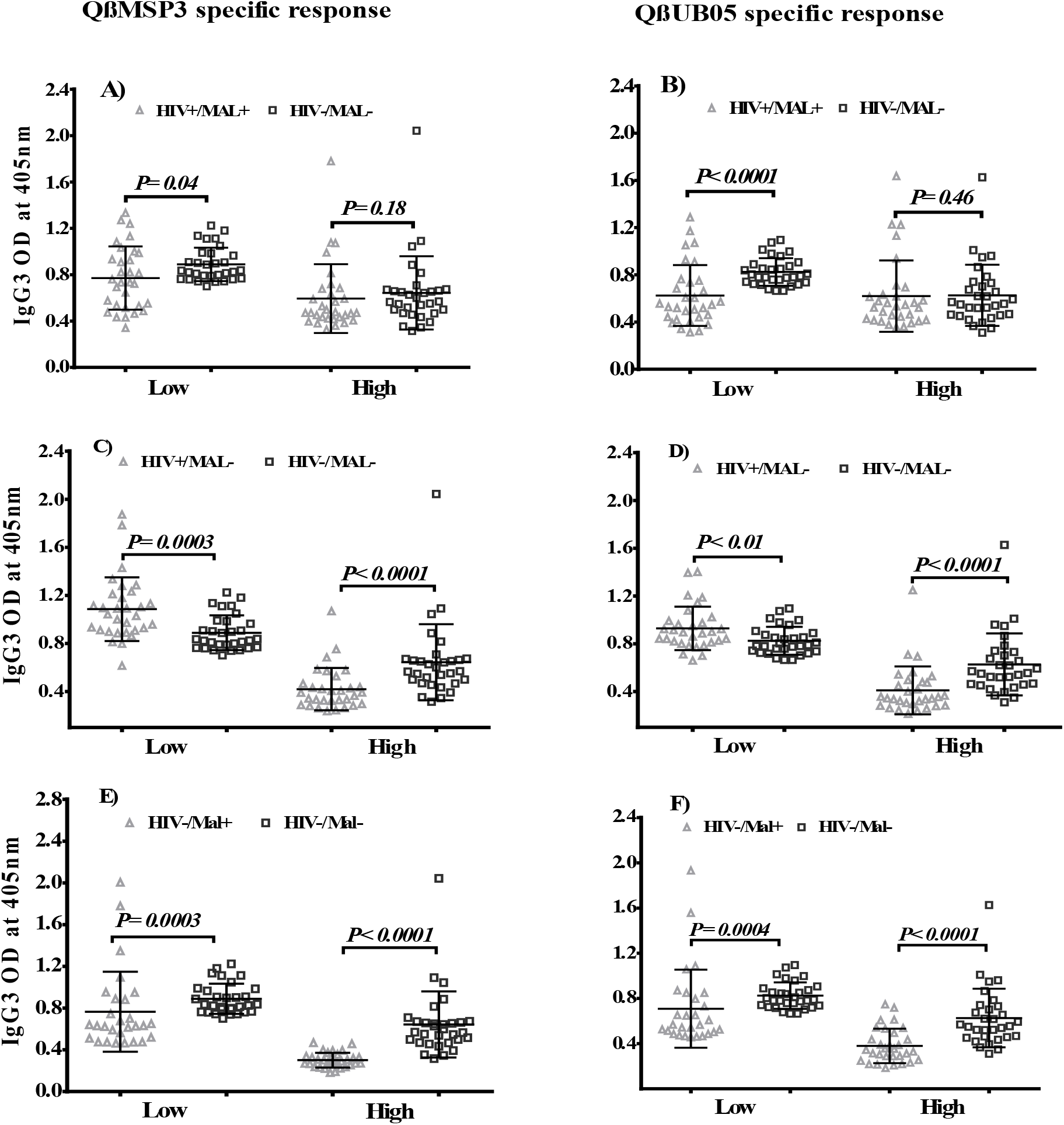
IgG3 specific response to the recombinant phages QβMSP3 and QβUB05 with respect to dual HIV-Malaria infection in individuals living in low and high malaria transmission areas. Comparison of IgG3 antibody responses between dual HIV-malaria infected and double negative individuals in low and high malaria transmission areas. IgG3 responses specific to the recombinant phages QβMSP3 (A) and QβUB05 (B) in low and high transmission areas. Effect of HIV infection on QβMSP3 (C) and QβUB05 (D) specific IgG3 responses in the two transmission areas. IgG3 responses specific to QβMSP3 (E) and QβUB05 (F) in individuals positive for *Plasmodium falciparum* in both transmission areas.

**Figure 5:**
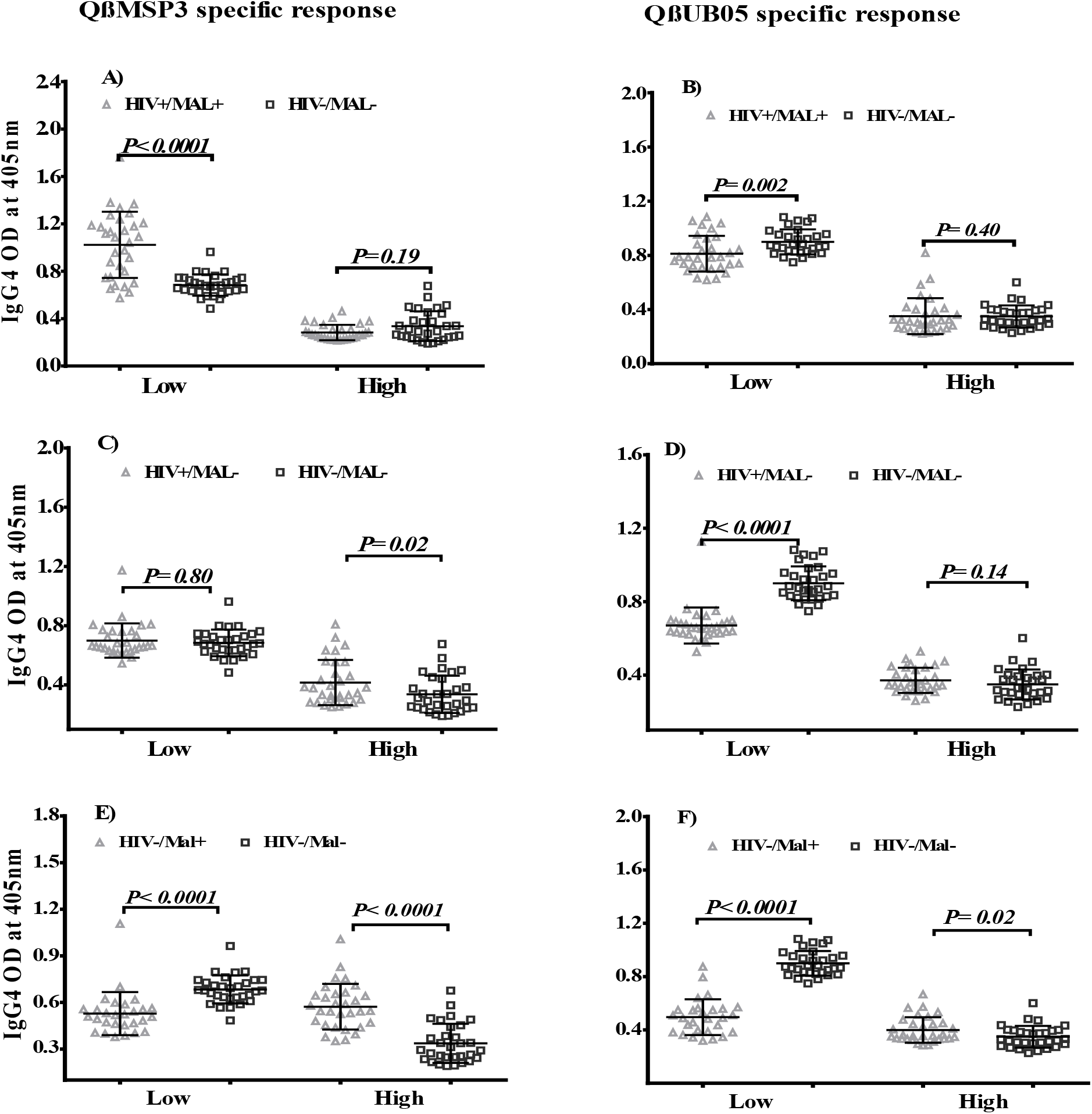
IgG4 response specific to the recombinant phages QβMSP3 and QβUB05 with respect to dual HIV-Malaria infection in individuals living in low and high malaria transmission areas. Comparison of IgG4 antibody responses between dual HIV-malaria infected and double negative individuals in low and high malaria transmission areas. IgG4 responses specific to the recombinant phages QβMSP3 (A) and QβUB05 (B) in low and high transmission areas. Effect of HIV infection on QβMSP3 (C) and QβUB05 (D) specific IgG4 responses in the two transmission areas. IgG responses specific to QβMSP3 (E) and QβUB05 (F) in individuals positive for *Plasmodium falciparum* in both transmission areas.

## Discussion

In this population based cross sectional study we profiled IgG and IgG subclass immune responses to QβUB05 and QβMSP3 in dual HIV-malaria infected people living in two areas of Cameroon differing in malaria transmission intensity. MSP3 and UB05 are asexual blood stage antigens known to be associated with naturally acquired immunity in individuals living in malaria endemic regions (18, 21, 23). However in populations living in malaria endemic regions parasite prevalence rate differ significantly between areas of low and high transmission (35). Since naturally acquired immunity to malaria is dependent upon the cumulative exposure in endemic regions low malaria transmission could limit exposure to parasites which has been linked to a waning antimalarial immunity and an increase in clinical episodes of malaria in adults (36–38). This in effect can modulate the profiles of target antigen specific IgG and IgG subclass responses that can be achieved especially when the individuals are co-infected with HIV which depletes the immune system. The impact of a high malaria transmission intensity in both dual HIV-malaria infected and double negative individuals was a significant increase in antigen specific IgG irrespective of the targeted antigen type. This might be in line with previous reports suggesting that high levels of antibodies to several blood stage antigens (4–7, 36) were a necessary component of protective immunity to malaria. Whereas this is probably correct for IgG antibody levels specific to QβUB05 and QβMSP3 in high malaria transmission areas the comparatively lower antibody levels in low transmission areas indicates that partial immunity to malaria could be waning in these areas due to a reduction in parasite antigen challenge (39) HIV infection resulted to a significant reduction in IgG antibody responses specific to QβMSP3 in both low (P=0.0001) and high (P=0.0001) malaria transmission areas. In contrast, IgG antibody responses specific to QβUB05 were not significantly affected by HIV infection (compares S1A&B). Infection with *Plasmodium falciparum* was associated with a significant increase in IgG antibodies specific for both antigens in the two transmission areas. This is especially true when dual HIV-malaria infected individuals are compared with participants infected with HIV alone. Here we demonstrated that even when there was HIV infection the impact of *Plasmodium falciparum* infection was a significant increase in IgG antibodies specific to MSP3 in both low (P=0.0001) and high (P=0.0001) malaria transmission areas (S1E&F).This indicates that asexual blood stage antigens are capable of modulating parasite antigen specific IgG antibody levels in both low and high malaria transmission areas. Such antibody levels are influenced by other factors including malaria and HIV infection.

When responses are examined qualitatively with respect to IgG subclasses and dual HIV-malaria infections differential outcomes are observed. In comparing IgG1 responses to the two antigens QβMSP3 specific IgG1 was associated with protection against malaria in low transmission areas in contrast to QβUB05 specific IgG1 responses which were relevant to protection in high malaria transmission areas. In this regards IgG1 responses specific to QβUB05 in high malaria transmission areas were significantly higher than responses to QβMSP3 both for dual HIV-malaria infected (P=0.004) and double negative (P=0.0001) individuals. This is probably in line with previous reports which associated increase UB05 specific antibody responses with recovery from clinical malaria (18). Similarly significantly higher IgG2 and IgG3 responses specific to both antigens were associated with protection in low transmission areas. This is in line with previous findings indicating that high levels of malaria parasite antigen specific IgG3 and IgG2 were relevant to protection against all forms of malaria (23, 40). This probably indicates that in malaria endemic regions novel recombinant phages such as QβUB05 could be more useful for detecting protective levels of antibodies implicated in the control of malaria in both low and high transmission areas. On the other hand the recombinant phage QβMSP3 would be more relevant for profiling IgG subclass responses in low malaria transmission areas.

The effect of dual HIV-malaria infection was a significant reduction in QβMSP3 specific IgG1 (P=0.0001), IgG2 (P=0.0001) and IgG3 (P=0.04) responses in low transmission areas. In low transmission areas this significant reduction of QβMSP3 specific IgG2 together with the cytophilic antibodies IgG1 and IgG3 in dual HIV-malaria infected people could diminish protection from *Plasmodium falciparum* infection and symptomatic illness. In the case of IgG subclass antibodies specific to QβUB05 such a reduction in dual HIV-malaria infected people might lead to increase morbidity and mortality to malaria in both low and high transmission areas. This is mainly because parasite antigen specific IgG1 and IgG3 are critical in monocyte mediated antibody-dependent cellular inhibition (ADCI) of *Plasmodium falciparum* which is responsible for killing asexual blood stages. Other reports have also shown that dual HIV-malaria infections escalate episodes of symptomatic malaria (41)or severe disease in both children and adults (42, 43). In endemic regions such individuals due to persistent parasitaemia could serve as reservoirs of malaria parasite thereby sustaining infection in both low and high transmission areas.

In low transmission areas dual HIV-malaria infection resulted to significantly higher QβMSP3 specific IgG4 responses which are probably the reason why IgG1 and IgG3 responses to QβMSP3 were significantly lower in this group. On the other hand, IgG4 responses to QβUB05 were significantly higher in double negative participants relative to dual HIV-malaria infected individuals. In high transmission areas IgG4 responses to both antigens were low which could explain why IgG antibody responses were comparatively high in this area. Since IgG4 is a noncytophilic IgG subclass which may block antibody mediated natural immunity to malaria (23, 32, 44–46) high IgG4 levels tended to be associated with lower IgG1 or IgG3 antibody levels. Overall the antibody responses to both antigens are relatively heterogeneous being certainly influenced by a number of factors including transmission area, ongoing *plasmodium falciparum* infection and coinfection with HIV. Never the less there was a clear indication that MSP3 specific IgG subclass responses were associated with protective immunity to malaria mainly in low transmission areas whilst similar responses to QβUB05 were related to protection in both low and high malaria transmission areas. This implies that QβMSP3 might not be suitable as a standalone vaccine in areas differing in transmission intensity. On the other hand antigenicity of UB05 most likely predicts immunity in both low and high transmission areas and could be used either alone or in combination with other antigens for vaccine studies in areas differing in transmission intensities. Thus understanding immune responses to QβUB05 and QβMSP3 could enable the development of efficacious vaccines or commensurate immunotherapeutic strategies suitable for areas differing in malaria transmission intensity.

**Table 1:** Study population characteristics.

| Variables | HIV-/Mal- (n=62) |  | HIV-/Mal+ (n= 59) |  | HIV+/Mal- (n= 62) |  | HIV+/Mal+ (n= 62) |  |
| --- | --- | --- | --- | --- | --- | --- | --- | --- |
| Gender | Male | Female | Male | Female | Male | Female | Male | Female |
| Participants (%) | 30 (48) | 32 (52) | 30 (51) | 29 (49) | 44 (71) | 18 (29) | 44 (71) | 18 (29) |
| Median Age (IQR) | 30 (22- 34.5) | 35.5 (22.75-48.75) | 32 (23–46.25) | 29 (26-45) | 46 (36.25-58.25) | 36 (30-46.25) | 38 (31 - 46) | 42 (33-52) |
| Median CD4 count (cell/mm3) | N/A |  | N/A |  | 480.6 (IQR : 270.5-635) |  | 459.6 (IQR : 241.0–594.8) |  |

N/A = Not Applicable, IQR = interquartile range, Mal+= malaria positive

## Acknowledgement

The authors are grateful to Professor Alain B Waffo for provinding the recombinant QβMSP3 and QβUB05 phages. We would like to thank the personnel of tunites techniques of CIRCB, Centre de Santé Catholique de Bikop and Mvogbesi Yaounde for their help in collecting the blood samples. Most importantly our gratitude goes to members of the CIRCB AFRODEC cohort for consenting to participate in this project.

